# The impact of functional correlations on task information coding

**DOI:** 10.1101/2022.11.23.517699

**Authors:** Takuya Ito, John D. Murray

## Abstract

State-dependent neural correlations can be understood from a neural coding framework. Noise correlations – trial-to-trial or moment-to-moment co-variability – can be interpreted only if the underlying signal correlation – similarity of task selectivity between pairs of neural units – is known. Despite many investigations in local spiking circuits, it remains unclear how this coding framework applies to large-scale brain networks. Here we investigated relationships between large-scale noise correlations and signal correlations in a multi-task human fMRI dataset. We found that task-state noise correlation changes (e.g., functional connectivity) did not typically change in the same direction as their underlying signal correlation (e.g., tuning similarity of two regions). This suggests that 1) trial-by-trial variability typically decreases between similarly tuned regions, and 2) stimulus-driven activity does not linearly superimpose atop the network’s background activity. Crucially, noise correlations that changed in the opposite direction as their signal correlation (i.e., anti-aligned correlations) improved information coding of these brain regions. In contrast, noise correlations that changed in the same direction (aligned noise correlations) as their signal correlation did not. Interestingly, these aligned noise correlations were primarily correlation increases, suggesting that most functional correlation increases across fMRI networks actually degrade information coding. These findings illustrate that state-dependent noise correlations shape information coding of functional brain networks, with interpretation of correlation changes requiring knowledge of underlying signal correlations.

## Introduction

Advances in functional brain imaging have enabled the investigation of the large-scale network organization of the human brain. In resting-state functional magnetic resonance imaging (fMRI), studies have found highly reliable and modular functional connectivity (FC) organization, which is measured through correlating the spontaneous fMRI activity of different brain regions (Power et al., 2011; Yeo et al., 2011). Related work has shown that this overall network organization persists across task states (Cole et al., 2014; Gonzalez-Castillo and Bandettini, 2017; Krienen et al., 2014), disease states (Spronk et al., 2021), and individuals (Gratton et al., 2018). Despite the appearance of a state– and trait-invariant network organization, there are reliable changes that occur to the network organization for specific networks or regions (Cole et al., 2014; Krienen et al., 2014; Shine et al., 2016). While some recent methodological efforts in human brain imaging have worked to disambiguate the sources of state-specific network correlation changes (Cole et al., 2016; Duff et al., 2018), the significance and interpretation of these correlation changes remain unclear.

In parallel, empirical and theoretical neurophysiological studies have established a rigorous statistical framework to study the properties of neural correlations and how they impact neural coding (Moreno-Bote et al., 2014; Panzeri et al., 2022; da Silveira and Berry, 2014). Critically, there are two forms of correlated activity that contain distinct sources of variance within neural data, yet provide complementary information about task coding: the signal correlation (SC) and the noise correlation (NC) (Cohen and Kohn, 2011). Intuitively, SC measures the tuning similarity between a pair of neural units. NC measures the temporal correlation during a task/stimulus (e.g., timepoints or trials) (Fig. 1c), capturing the dynamic interaction of two units. (Note that the terms SC and NC were originally defined through the lens of information theory, where “signal” corresponds to the mean across responses, and “noise” corresponds to the variance across responses (MacKay, 2003). In the context of prior fMRI literature, these SCs and NCs are statistically equivalent to across-task co-activations and FC, respectively (Cole et al., 2019).) Under the theoretical neural coding framework, studies have suggested that the effect a NC has on task coding depends on how well it aligns i.e., shares the same sign) with the underlying SC of those two units (Moreno-Bote et al., 2014; Panzeri et al., 2022; da Silveira and Berry, 2014). In particular, the signal-noise angle – the difference in the directions (sign and magnitude) of the SC and NC – determines the information coding properties of a neural population (Panzeri et al., 2022). This is because if an NC aligns with its SC, this would interfere with the coding direction of these two units. While the theoretical account of SC/NC was developed to account for empirical phenomena observed at the level of neuron pairs during the presentation of fine-grained sensory stimuli (Cohen and Kohn, 2011; Moreno-Bote et al., 2014; da Silveira and Berry, 2014) and local fMRI voxels (Zhang et al., 2020; Cheng et al., 2022; van Bergen and Jehee, 2018), the statistical principles are generic to account for neural data across a wide range of spatial and cognitive scales, including large-scale brain networks. Thus, we sought to investigate whether the SC/NC coding principles apply to the level of large-scale brain regions. A successful demonstration of SC/NC coding principles at the level of human functional brain networks would bridge the vast literature of FC analyses prevalent in the human functional connectomics literature with the rich neural coding framework developed in theoretical neuroscience.

**Figure 1.**
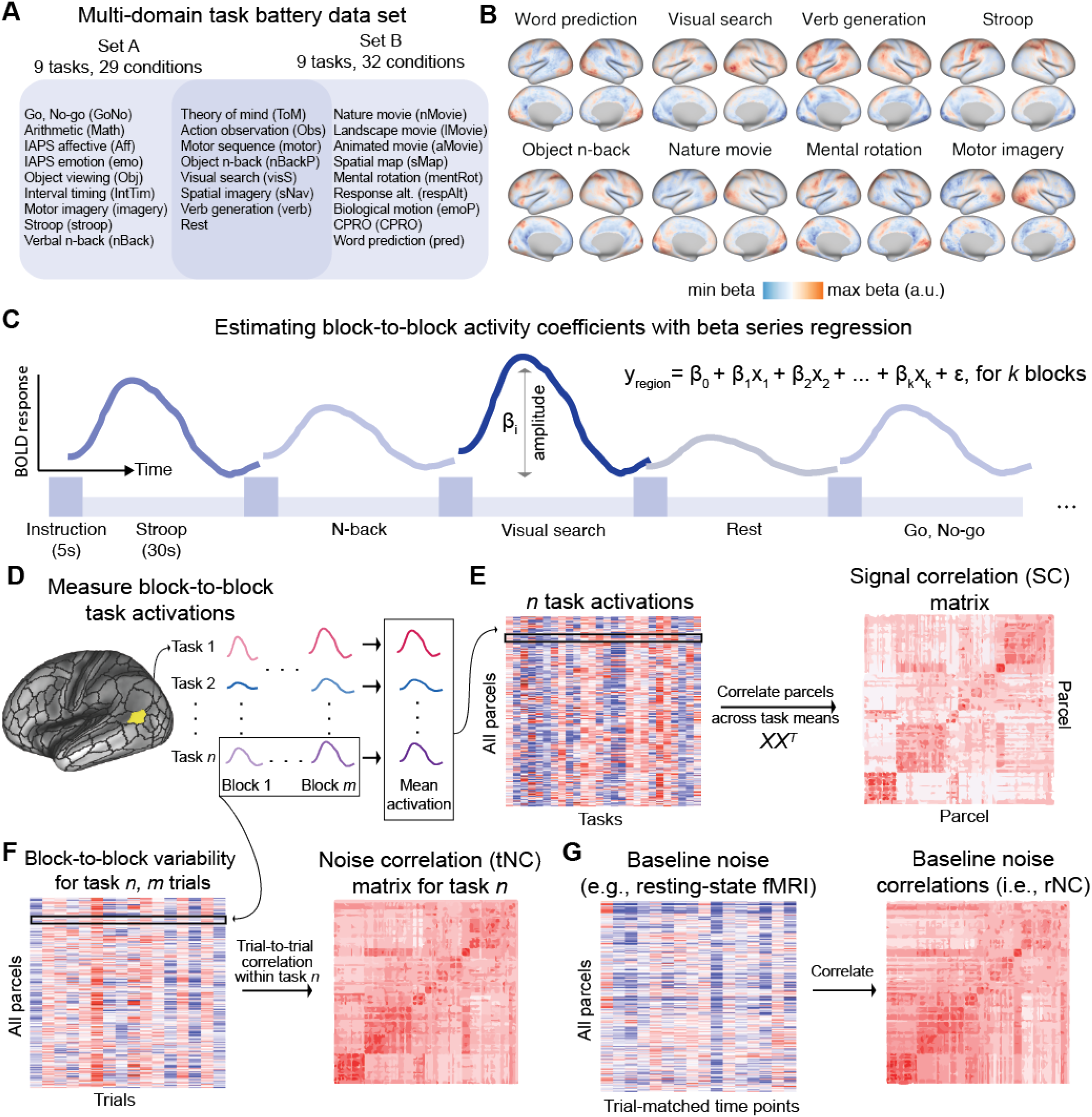
We used a multi-task dataset to capture the large-scale SC and NC organization in human functional brain networks. a) The MDTB dataset with 26 unique tasks (King et al., 2019). b) Cortical activation maps for eight example tasks. c) Block-wise activation estimates were obtained using a beta series regression approach, where each task block was modeled independently in a linear regression model (Rissman et al., 2004). d) SCs and NCs in large-scale fMRI data are estimated from orthogonal timeseries sources. We estimate the trial-to-trial task activation amplitude in fMRI data for each region, and for all tasks. e) To estimate the SC matrix, we compute the correlation between all pairs of brain parcels using the cross-trial mean activation of many tasks. f) In contrast, tNC matrices for a given task are computed as the correlation of trial-to-trial variability between pairs of parcels within a single task. g) The tNC can be compared to the well-studied baseline rNC using resting-state fMRI activity. SCs and NCs are computed for each participant separately, and then averaged to produce a group-level matrix estimate.

SCs capture the similarity of task tuning between two neural units (neurons or brain regions). At the level of single neurons, this typically captures the tuning curve similarity of fine-grained sensory stimuli, such as the orientation of visual gratings (Cohen and Maunsell, 2009). While the impact of these correlations on task coding have been widely investigated in local spiking circuits, it remains unclear how this coding framework applies to large-scale brain networks. This is largely due to the fact that the types of fine-grained tuning curves (i.e., orientation gratings) captured in prior neurophysiology studies are generally inaccessible at the level of large-scale fMRI brain networks. Instead, large-scale fMRI brain networks have been previously shown to be selective to broader cognitive tuning curves, such as different cognitive tasks (Yeo et al., 2015; Smith et al., 2009). Thus, we leverage a multi-task dataset that spans diverse cognitive domains to characterize the SC and NC organization across distributed functional brain networks. This approach plays to the strengths of fMRI, while allowing us to extend the prior theoretical neural coding framework to large-scale functional brain networks.

Here we characterize the organization of SC and NC in large-scale human brain networks, and assess their coding properties across a wide range of cognitive tasks. While prior human neuroimaging studies primarily viewed correlations (i.e., FC) through the lens of dynamic communication (for a review, see Gonzalez-Castillo and Bandettini (2017)), we test whether FC can be interpreted through the lens of information coding. (Since FC and NC are statistically equivalent, we use them interchangeably in this study; see Table 1.) First, we extend the notion of SC from the correlation of visual tuning curves to a wide variety of cognitive tasks (i.e., *cognitive tuning curves*). We compared the organization of SC to the well-established resting-state NC (rNC) organization of human cortex, finding that SC reflected a more modular and segregated network organization than rNC. Next, we built a statistical model of NC which demonstrated that, under the assumption that observed NC is driven by a linear combination of internal neural and external task sources, NCs should exclusively change in the direction of their underlying SC (i.e., positive increases in NC should be observed when the SC is positive). In contrast to this assumption, we found that NC changes do not typically align with the underlying SC in empirical fMRI data. Instead, a majority of NCs changed in the direction that was opposite to the SC. To understand the functional relevance of these NC changes, we leveraged the hypothesis from theoretical neuroscience that the alignment of the signal and noise correlations impacts the fidelity of task information coding. Indeed, we found that the signal-noise relationship is predictive of the fidelity of task coding in large-scale brain networks. These results shed light on the relationship between neural correlations and information coding, placing fMRI functional connectomics within the broader neural coding framework.

**Table 1.** Table of definitions and abbreviations. Distinct terms used in different subfields within neuroscience are often computed identically and have converging interpretations. For example, noise correlations and functional connectivity are computed in a statistically identical manner, and aim to capture a similar empirical phenomenon: interaction of neural units.

| Name | Abbreviation | Interpretation |
| --- | --- | --- |
| Signal correlation | SC | Similarity of task selectivity |
| Cross-task co-activations | n/a | *Across-task correlation of mean activations* |
| Noise correlation | NC | Neural/dynamic interactions |
| Functional connectivity | FC | *Correlation of ongoing activity* |
| Resting-state noise correlations | rNC | Spontaneous interactions/correlations |
| Resting-state functional connectivity | rFC | *Correlation of ongoing spontaneous variance* |
| Task-state noise correlations | tNC | Task-state interactions/correlations |
| Task-state functional connectivity | tFC | *Correlation of ongoing task-driven variance* |
| Noise correlation change | ΔNC | Task versus rest/baseline correlation change |
| Task vs rest FC change | ΔFC | *Difference between task and rest correlations* |

## Results

### Estimating multi-task SCs and NCs in human functional brain networks

We first characterized multi-task SC and NC across all pairs of parcels (see Table 1 for definitions). We used the publicly available multi-domain task battery dataset collected by King and colleagues (King et al., 2019). Briefly, the multi-domain task battery dataset contains 26 cognitive tasks per participant (Fig. 1a). Tasks were interleaved across blocks, where each block was preceded by a 5s instruction screen, followed by a 30s block. For our analyses, we modeled the mean activity of each block separately for every brain region (parcel) in the Glasser atlas (Glasser et al., 2016) using a beta series regression (Rissman et al., 2004) (Fig. 1c).

To compute the SC between all pairs of parcels, we first computed the mean activation across blocks for each task separately (Fig. 1d). This yielded a 360 parcel by 26 task matrix, from which we computed the SC matrix (Fig. 1e). NC was calculated using the cross-block variability for every parcel, which is a distinct statistical property to the cross-block mean. (Note every task had the same number of blocks.) We calculated the NC between all pairs of brain regions, and across all tasks (Fig. 1f). To get a task-state NC (tNC) matrix, we averaged the NC across all tasks (excluding the resting-state condition). Resting-state blocks were also interleaved throughout the experimental design. To maintain consistency with how resting-state and task-state NC were computed, resting-state NC (rNC) was computed in an identical manner to tNC (i.e., using a beta series regression) (Fig. 1g). Note that the across-block rNC matrix estimated here is quantitatively similar to the more common rNC that is computed across timepoints in the human neuroimaging literature (Supplementary Fig. 1). Conceptually, the approaches are equivalent in that NCs capture the variability across task responses, and SCs capture the mean across task responses. Here we opt for cross-block analysis, since it enables the characterization of task coding for each block, rather than across timepoints. There were no statistically significant differences in movement (i.e., framewise displacement) during rest and task blocks (Supplementary Fig. 2). Furthermore, we ensured both SC, rNC, and tNC matrices were highly reliable by measuring the reliability across subject splits, with a reliability of 0.81 or greater (Supplementary Fig. 3e).

### SCs reveal a highly modular and segregated network organization

We characterized the SC matrix in the context of the well-known rNC matrix. Prior work in rNC studies revealed a modular organization of functional brain networks (Fig. 2a,c) (Ji et al., 2019; Power et al., 2011; Yeo et al., 2011). These functional network divisions were identified using clustering and community detection algorithms on resting-state NC matrices. To evaluate how SC was related to this modular network organization, we computed the modularity and segregation of SC with respect to the previously-defined resting-state network partitions (Fig. 2a). Modularity and segregation are related statistics that measure the strength of nodes within a network relative to the between-network connection strength (see Methods). Surprisingly, while the network partitions were optimized to maximize modularity from resting-state NC data, we found that SC had both higher modularity and segregation than resting-state NC (Fig. 2c). This suggests that SC can recapitulate the well-known functional subdivisions of cortex that are extracted from resting-state NC.

**Figure 2.**
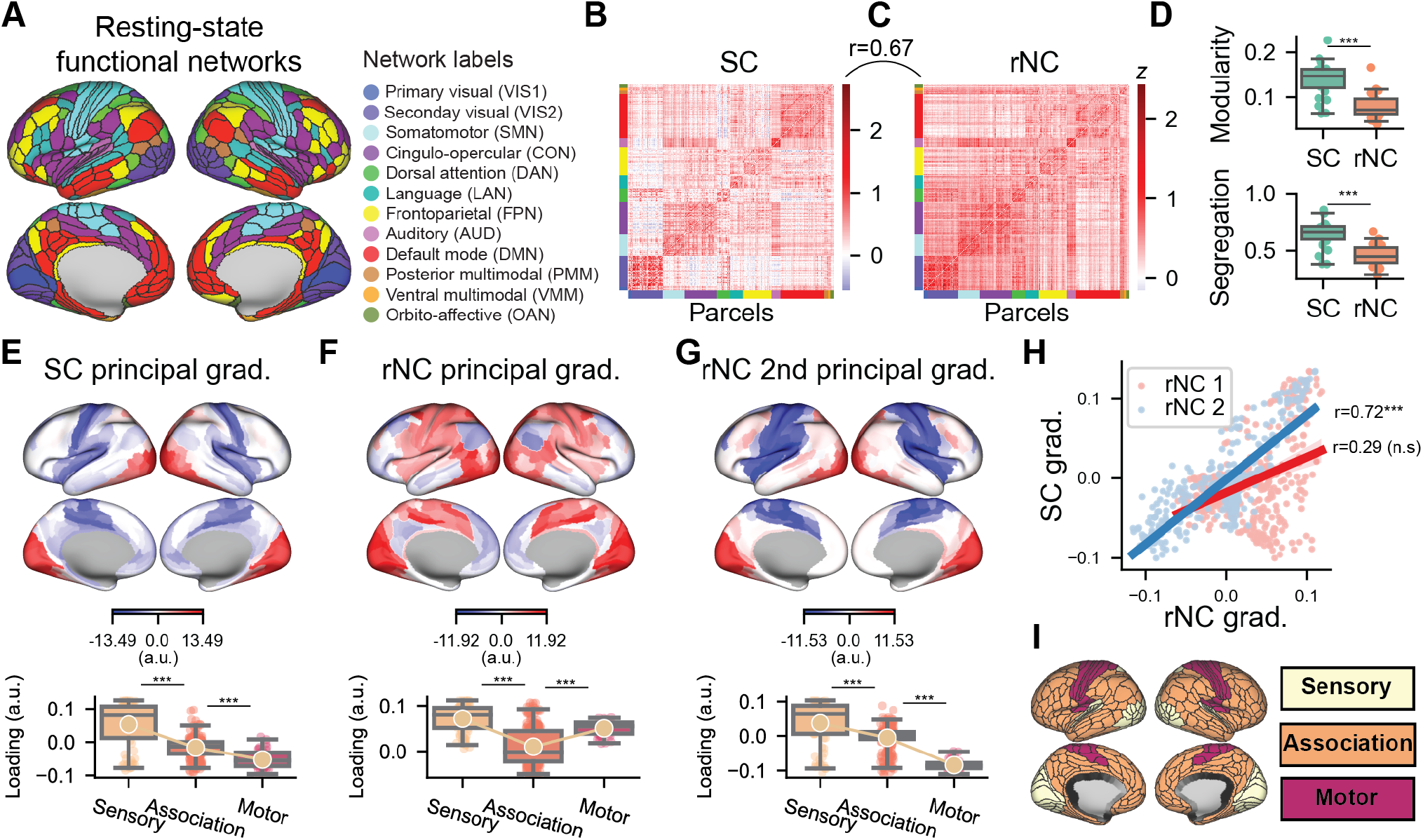
Comparing the SC matrix to the well-studied rNC matrix. a) We used the Glasser parcellation with 360 cortical parcels. Parcels were partitioned into 12 functional networks (Ji et al., 2019). b) The SC matrix, which captures the task tuning similarity between pairs of brain regions. c) The rNC matrix. d) Modularity and segregation (using the Ji et al. partition) of the SC and rNC matrices. e) Top: The first principal component of the SC matrix aligns along a sensory-association-motor gradient. Bottom: Average loading projected onto three cortical systems. f) The first principal component of the rNC is organized along the unimodal-transmodal (i.e., sensorimotor-association) hierarchy. g) The second principal component of the rNC matrix also aligns along a sensory-to-motor gradient, and is h) highly correlated with the SC principal gradient. i) Sensory, association, and motor systems projected onto the cortex. A full comparison of the first three gradients of the SC and rNC can be found in Supplementary Fig. 4. Box plot bounds define the first and third quartiles (across participants), box whiskers indicate the 95% confidence interval, and the center line indicates the median.

Note, however, that empirically estimating SC is a challenging problem. This is because when estimating within-subject SC, undesirable noise sources that are idiosyncratic with that individual and/or fMRI session may seep into the analysis, potentially confounding the ‘signal’ with subject-specific noise. One alternative to the naive approach of estimating SC within an individual, is to compute the inter-subject SC (Kim et al., 2018). This estimates the group-level SC using the task signal vector of one subject, with the task signal vector of another subject (or group of subjects). This isolates the task-driven signals, while ensuring that the individual-specific noise does not contaminate SC. When computing inter-subject SC, we found strong correspondence with within-subject SC and inter-subject SC (r=0.81), providing validation that SC provides reliable information about large-scale cognitive tuning curves (Supplementary Fig. 3.

### Cortical SCs are organized along a gradient of functional specialization

Complementing network analyses of SC and NC, gradient analysis offers a way to capture the greatest axes of variation of the entire SC and NC matrices (Margulies et al., 2016; Huntenburg et al., 2018). Gradient organization is computed by performing dimensionality reduction on the SC (or NC) matrices (e.g., a principal component analysis), and is complementary to network partitions as they exhibit smooth loadings/partitions, rather than “hard” or non-overlapping networks (Huntenburg et al., 2018). The first gradient of the rNC matrix, which is equivalent to its first principal component, is the well-documented sensorimotor-association (or unimodal-transmodal) hierarchy that was first described by Mesulam (Mesulam, 1998), and subsequently identified in fMRI data (Margulies et al., 2016) (Fig. 2f). This unimodal-transmodal gradient is related to both transcriptomic variation (Burt et al., 2018) and myelination content, which is captured in the T1w/T2w contrast map (Glasser and Van Essen, 2011) (26.1% variance explained; Supplementary Fig. 4h). However, gradient analysis of the SC matrix revealed a gradient of functional specialization, from sensory-association-motor areas (23.0% variance explained; Fig. 2e; see Supplementary Fig. 4 for additional details.). Critically, when grouping together cortical systems into sensory, association, and motor systems (Fig. 2i) – systems that are functionally distinct/specialized from each other – we found a monotonic relationship between these systems and their gradient loading. This is consistent with a gradient of functional specialization, where sensory and motor regions are defined by distinct functions, while association regions integrate the two (Ito and Murray, 2023). Moreover, this sensory-to-motor SC gradient was significantly associated with the 2nd principal gradient of rNC (rank r=0.72, non-parametric p<0.001) (Fig. 2h). By comparison, the SC gradient was not correlated with the typical unimodal-transmodal gradient (i.e., the 1st rNC principal gradient; rank r=0.29, p=0.16). Note that the relative difference in variance explained between the first and second gradients was larger for the SC matrix (Δ3.68% for the rNC matrix, and Δ5.45% in the SC matrix; see Supplementary Fig. 4h, suggesting that the re-ordering of the gradients in the SC matrix is not due to marginal changes in the amount of variance explained per gradient. Together, these results illustrate that while SC preserves the overall functional brain network organization, it reveals a more cognitively specialized organization that more clearly delineates functionally specialized regions. This is consistent with the notion that SCs capture task selectivity similarities between brain regions.

### A linear model of state-specific SC and NC changes

A brain region’s functional specificity emerges from its pattern of connectivity, i.e., its connectivity fingerprint (Passingham et al., 2002; Mars et al., 2018). Thus, two regions with similar functions or tuning curves (i.e., high positive SC) are likely to have high amounts of shared spontaneous activity (due to strong functional connections; i.e., high positive rNC). We verified this in our empirical data, finding that the SC matrix had overall strong correspondence with rNC (rank r=0.67, p*<*0.0001; Fig. 2b,c). However, how should stimulus-driven activity interact with spontaneous activity? To gain intuition on the interaction between stimulus-driven and spontaneous activity, we constructed a statistical model to simulate how state-specific NC emerges from an anatomically-constrained network model with linear dynamics.

We constructed a linear statistical model with 360 units and 10 networks (36 units per network). A unit’s activity was determined by the algebraic sum between shared baseline activity (shared between units in the same network), stimulus-driven noise, private noise (for each unit separately), and a globally shared signal (inducing positive correlations between all units; see Methods) (Fig. 3a,b). We found that this model produced positively correlated activity amongst all pairs of units (mimicking our empirical data), with greater correlation between units within the same network (i.e., shared connections; Fig. 3c,d). Critically, when including an additional stimulus-driven component, this primarily increased the magnitude of correlation primarily between strongly connected units (Fig. 3e). This model indicates that under the assumption of linearity, neural units that receive shared input drive should increase the magnitude of their correlated activity. In other words, with the addition of a new stimulus-drive, the state-dependent ΔNC should align with the strength of the underlying SC.

**Figure 3.**
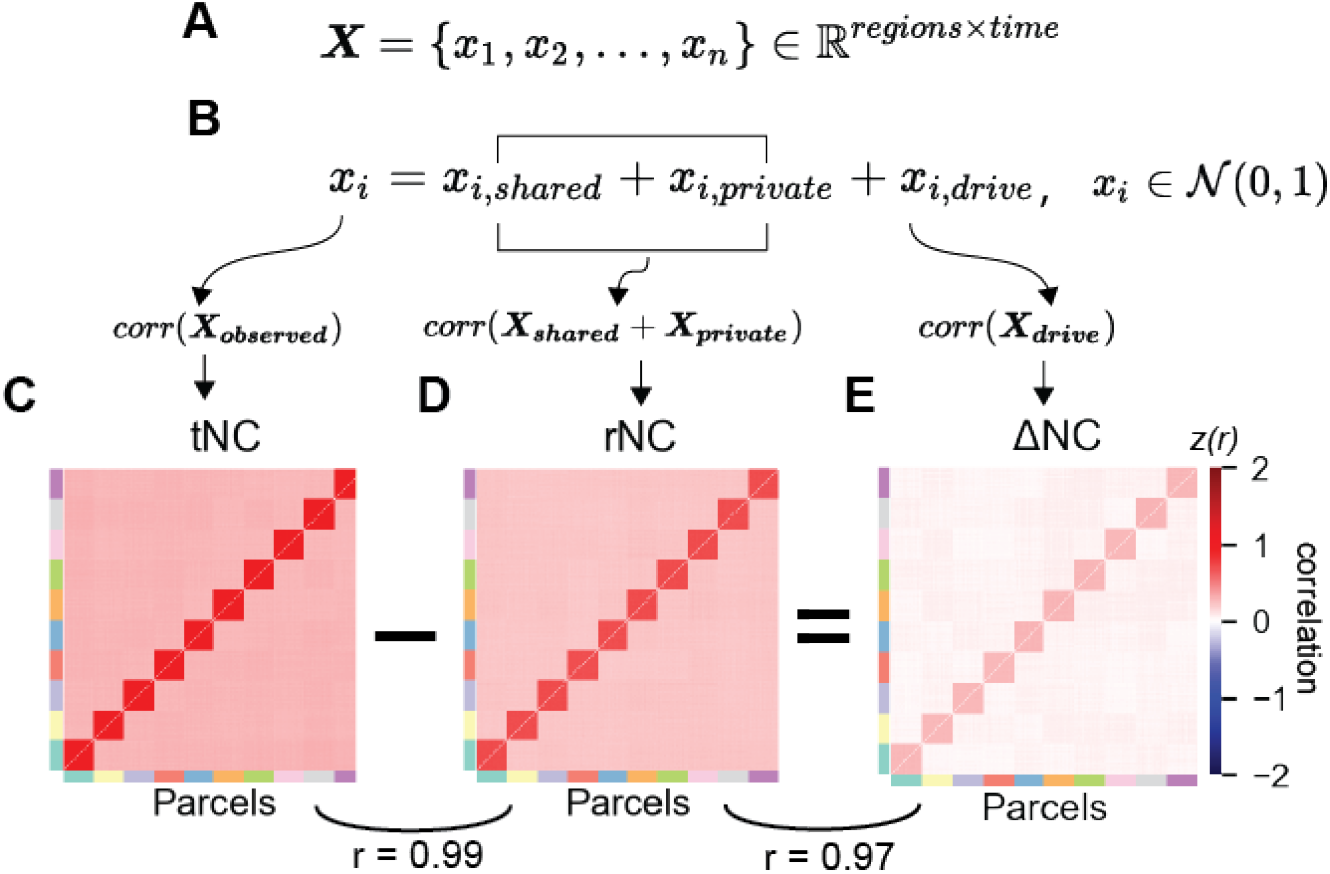
A linear model predicts neural dynamics during tasks can be decomposed into separable components. a) We constructed a simple network model with *n* = 360 units and 10 connected networks. b) To obtain the neural dynamics of a single brain region *x*_*i*_, linear dynamics were superimposed atop anatomical connectivity with a node’s private dynamics (*x*_*i*,*private*_), a network’s shared dynamics (*x*_*i*,*shared*_), and task dynamics (*x*_*i*,*drive*_). (All *x*_*i*_ sampled from *N* (0, 1).) Under these simple assumptions, the difference between the c) observed tNC and the d) baseline rNC yields the e) stimulus-driven component of correlated activity (i.e., ΔNC) in empirical (or simulated data).

### NC changes do not typically align with the underlying SC in empirical data

The statistical model provided an intuition of what should be expected if spontaneous and task-driven variance linearly interact. We next sought to characterize the relationship between SC and state-related NCs in empirical data. We characterized the rNC, task-state NC, and the ΔNC between the two (Fig. 4a-c). Consistent with prior work (Ito et al., 2020), we found that the overall change in correlation was dominated by correlation decreases. We computed the signal-noise differential matrix, which we defined as the Hadamard (element-wise) matrix multiplication of the SC matrix (Fig. 4d) with the ΔNC matrix (Fig. 4c). Note that we calculated the signal-noise differential matrix using the ΔNC matrix since we wanted to understand the impact of state-dependent changes in NC relative to ongoing spontaneous activity. Contrary to the statistical model and other studies arguing that NC dynamics are linear (Nozari et al., 2020), we found that most state-related ΔNC did not align with its underlying SC (aligned ΔNC pairs=42.99%; anti-aligned ΔNC pairs=56.74%; Fig. 4f). This suggests that the majority state-related ΔNC changes cannot be explained by the linear superposition of stimulus-driven and spontaneous activity. This finding is also consistent with prior reports indicating that there are a combination of multiplicative and additive fluctuations in ongoing local variability in both rodent electrophysiology (Lin et al., 2015), non-human primate electrophysiology (Churchland et al., 2010), and in human fMRI (He, 2013).

**Figure 4.**
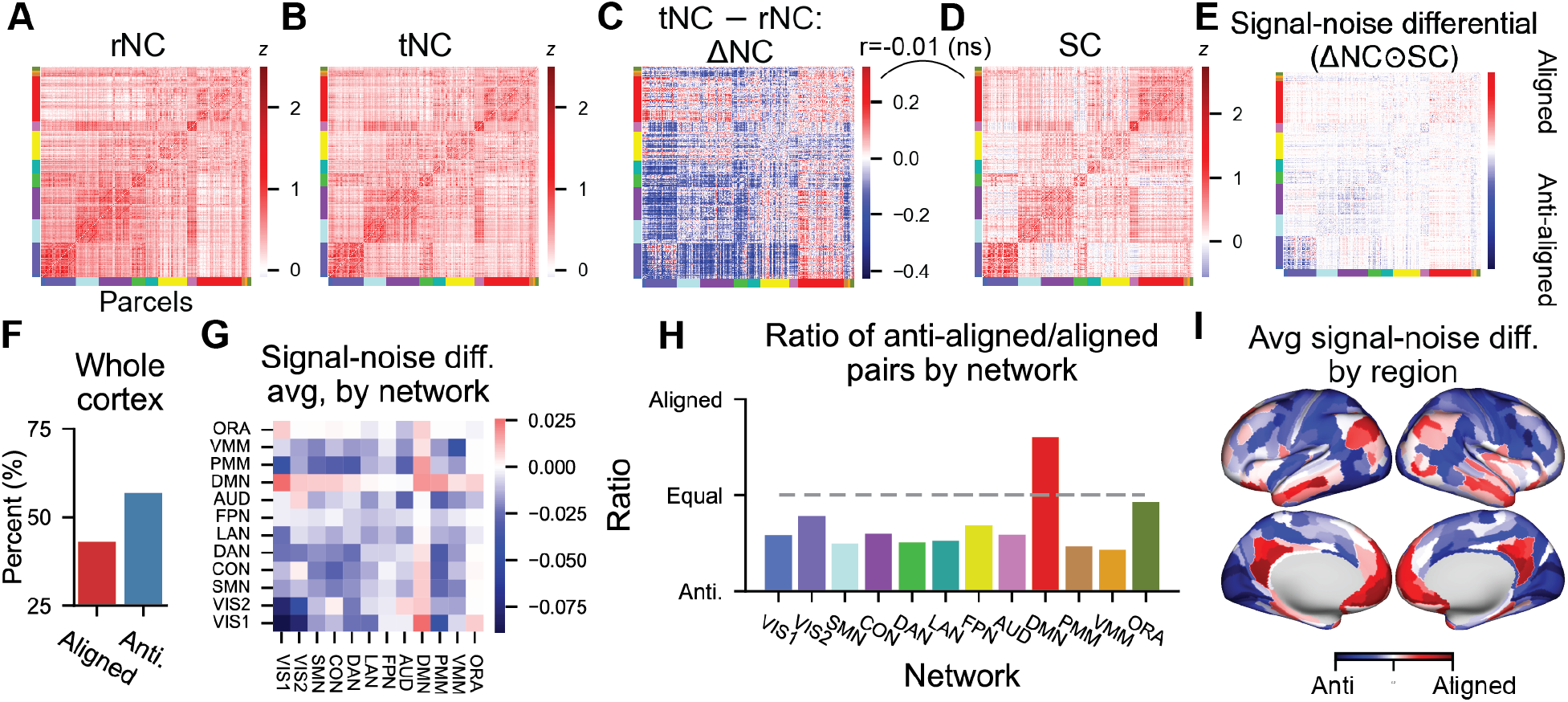
Disambiguating SC and state-dependent ΔNCs in functional brain networks using the signal-noise differential matrix. a) The rNC and b) tNC matrix. c) The tNC vs rNC matrix exhibits widespread correlation reductions. d) SC matrix, which reflects the task encoding similarity between pairs of regions. e) The signal-noise differential matrix can be obtained by computing the Hadamard product (element-wise multiplication) of the SC and the ΔNC matrix. The signal-noise differential matrix therefore reflects whether the ΔNC between a pair of regions reflects a change that is aligned (positive) or anti-aligned (negative) with its underlying SC. f) Percentage of aligned and anti-aligned signal-noise angle pairs across all cortical pairs. g) The signal-noise differential matrix averaged by network. h) Percent of aligned vs. anti-aligned NCs by each functional network. DMN is the only network that contains more aligned than anti-aligned NC changes. i) The average of the signal-noise differential matrix for each region (i.e., averaging across columns for each row in panel e.

We next characterized the network organization of aligned and anti-aligned ΔNCs. While the majority of networks were dominated by anti-aligned ΔNCs, the Default Mode Network (DMN) was instead dominated by aligned ΔNC pairs (Fig. 4g-i). The DMN, which primarily consists of the medial prefrontal cortex and posterior cingulate, has previously been shown to suppress its activity during task performance (Dosenbach et al., 2007; Raichle et al., 2001). Prior work characterizing the impact of NCs on neural coding suggest that an aligned signal-noise differential inhibits information coding due to the interference of the correlated noise along the coding (signal) axis (Panzeri et al., 2022). This predicts that the ΔNC increases associated with the DMN (Fig. 4c) may inhibit the coding of task-related information. In what follows, we provide a theoretical intuition of why an aligned signal-noise angle inhibits information coding, and directly test out this theory in fMRI data.

### Interpreting ΔNC through a neural coding framework

There is a rich history in neuroscience of investigating correlated neural activity through the lens of information coding (Johnson, 1980; Abbott and Dayan, 1999; Averbeck et al., 2006; Cohen and Kohn, 2011; Kohn et al., 2016; Moreno-Bote et al., 2014; Panzeri et al., 2022; da Silveira and Berry, 2014). Recent theoretical work suggested that the impact of the NC on information coding critically depends on the signs of the SC and NC (Moreno-Bote et al., 2014; da Silveira and Berry, 2014). This intuition can be geometrically described in terms of the signal-noise angle (Panzeri et al., 2022). The signal axis describes the direction of maximal covariance of the mean activity across many tasks/stimuli between a pair of neural units (Fig. 5a). The noise axis describes the direction of maximum noise covariance. That is, covariance across repeated instances (e.g., trials or blocks) of the same task/stimuli (5b-d). Thus, the signal-noise angle describes the angle between these two directions, and reflects whether the NC is information-enhancing (orthogonal to SC) or information-limiting (aligned to SC). However, this initially proposed framework only considers the overall magnitude of the NC, neglecting the impact of spontaneous rNC, which can be used as a baseline. However, prior work in human neuroimaging has shown that the spontaneous correlations estimated during resting-state fMRI are stable and informative (non-zero), reflecting an intrinsic network organization (Gratton et al., 2018). Thus, investigating the impact of NCs on information coding relative to baseline would shed light on how the brain dynamically reconfigures to support information-enhanced or information-limiting coding between pairs of brain regions.

**Figure 5.**
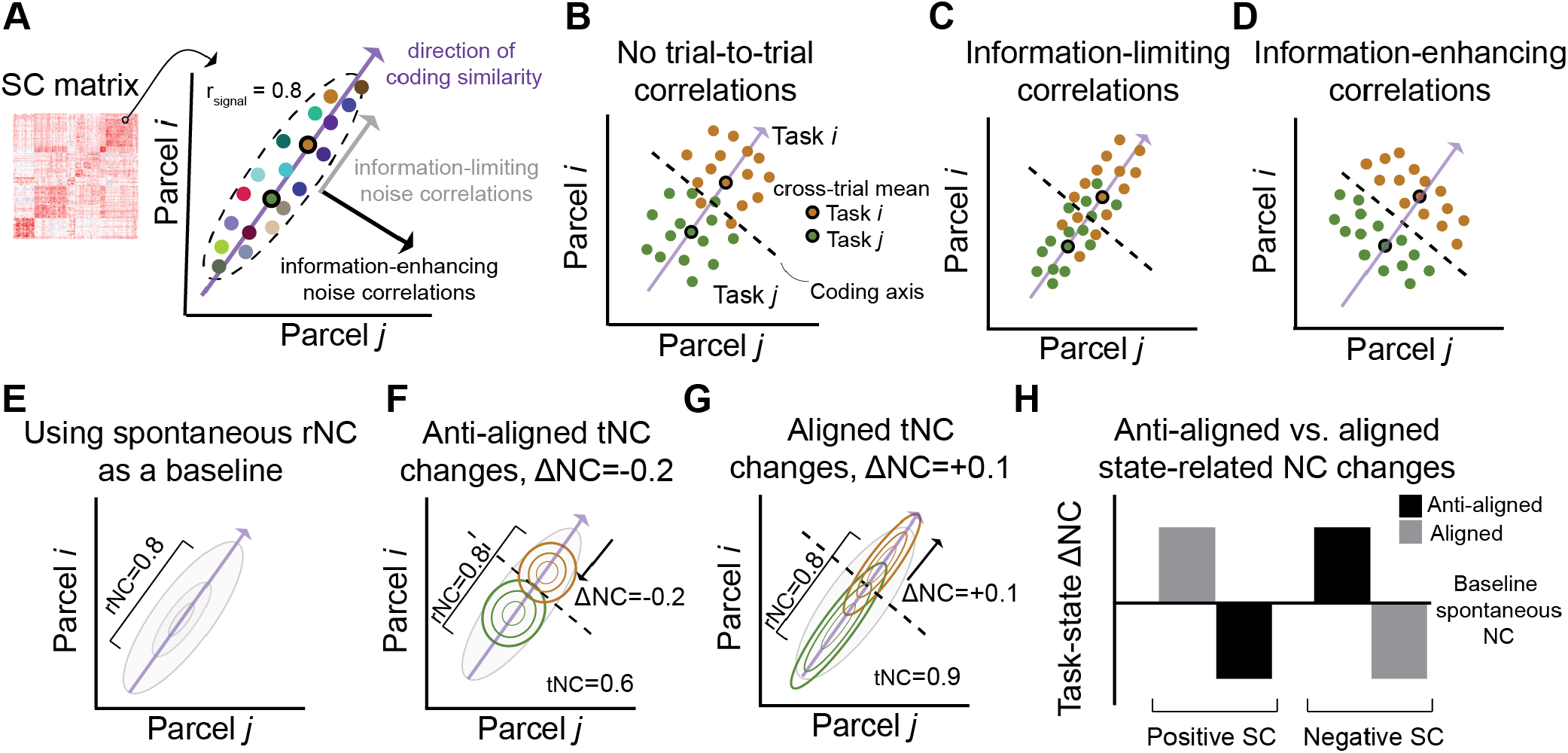
Interpreting ΔNC from an information coding framework. a) For a pair of brain regions, the SC captures the direction of maximal covariance of the mean activity across many tasks (each colored dot represents the mean activity of a distinct task). tNCs, on the other hand, capture within-task covariability (across events). b) An example of weak (or no) tNC for two tasks. c) Prior theories in the neural coding effects of tNC posit that correlations in the same direction as the underlying SC are information limiting. This is because the activity becomes more difficult for a linear decoder to distinguish between the two task conditions. d) In contrast, tNCs that are in the orthogonal direction as the underlying SC are information enhancing, since the trial-to-trial activity becomes more easily decodable by a linear classifier. e-g) We modify prior theories to assess how the task-state reconfiguration of tNC impacts information coding relative to the e) baseline rNC estimate. This modification involves estimating the ΔNC (tNC *−* rNC). f) An anti-aligned ΔNC, whereby the ΔNC is the opposite sign of the SC, thereby reducing noise interference along the SC axis. g) An aligned ΔNC, whereby the ΔNC is the same sign of the SC, thereby increasing noise interference along the SC axis. h) ΔNCs are putatively information-limiting or information-enhancing based on how the ΔNCs are aligned or anti-aligned with the underlying SC.

To assess the reconfiguration of NCs from a baseline state (i.e., rNC) to a task state (i.e., tNC), we made several modifications to prior theories. First, we measured the rNC to establish a baseline (Fig. 5e). Next, we measured the tNC, and computed the ΔNC (tNC *−* rNC) (Fig. 5f,g). If the ΔNC was of the same sign as the underlying SC (i.e., an aligned ΔNC; Fig. 5g), this would suggest that the brain dynamically reconfigured such the tNC would interfere the coding axis. In contrast, if the ΔNC was the opposite sign as the the underlying SC (i.e., an anti-aligned ΔNC; Fig. 5f), then we would infer that the brain dynamically reconfigures the NC such that the tNC minimizes interference along the SC axis relative to baseline. Therefore, the product of the SC and ΔNC (Fig. 5h) – which we define as the signal-noise differential – serves as a useful estimate to capture how ΔNC impacts information coding.

Though prior work in human neuroimaging has reported more prevalent negative correlations in the rNC matrix, these negative correlations are introduced through a preprocessing technique known as global signal regression (Murphy et al., 2009). Global signal regression artefactually reduces the mean (across the entire brain) NC to 0, making it difficult to directly compare the impact of magnitude differences across rest and task states. However, here we derive rNC and tNC from the same imaging sessions (where rest is interleaved with task), therefore ensuring that differences in NC values cannot be due to different baselines across different imaging runs. This ensures that the comparison of tNC and rNC magnitudes are interpretable. We next test the hypothesis that the relationship between SCs and ΔNCs impact task information coding in empirical fMRI brain networks.

### The signal-noise differential determines the impact of NCs on task information decoding

Theoretical work suggests that the signal-noise differential determines how easily task information can be decoded from a set of neural units. To test this empirically, we began by identifying sets of brain regions with entirely aligned or anti-aligned signal-noise differentials (i.e., Fig. 4e). We leveraged a technique from network science – clique identification – to identify groups of brain regions with exclusively aligned or anti-aligned signal-noise differentials (Palla et al., 2005) (Fig. 6a). Identifying cliques of either aligned or anti-aligned ΔNCs ensured that all brain regions would either have putatively information-limiting or information-enhancing correlations with each other.

**Figure 6.**
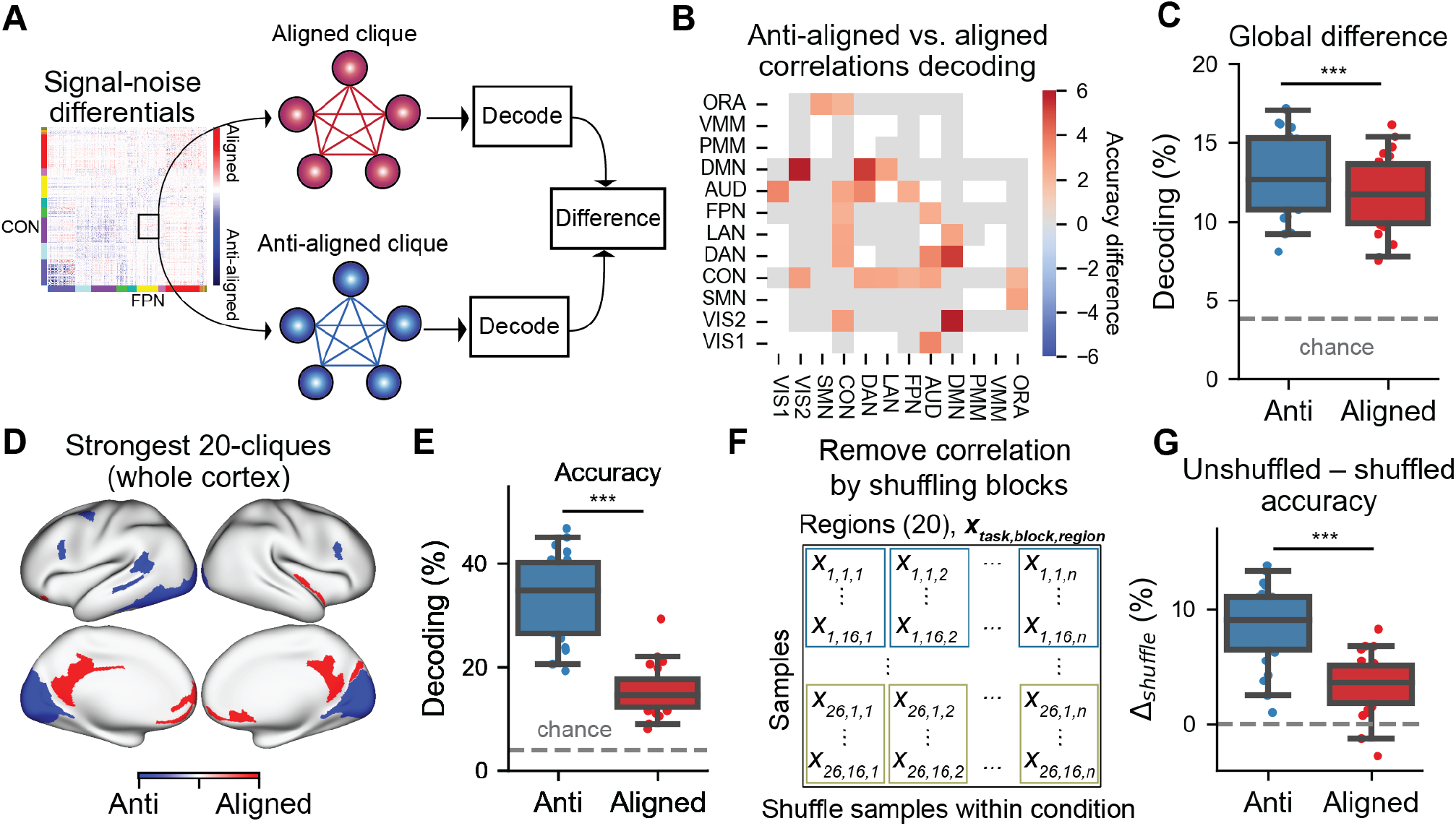
Brain regions with anti-aligned ΔNCs have improved information decoding over aligned ΔNCs. a) We identified network-matched sets of aligned or anti-aligned ΔNCs by identifying cliques (sub-networks of entirely aligned or anti-aligned ΔNCs). b) For each pair of networks, we found that sets of brain regions with anti-aligned ΔNCs had significantly higher multi-task decoding performance than brain regions with aligned ΔNCs for specific network pairs (FWE-corrected). Note that matrix elements colored in gray had no significant difference. Elements in white were not testable (due to non-existence of aligned and/or anti-aligned cliques). c) We computed the average difference for every matrix element in panel b) for anti-aligned versus aligned cliques, finding that on average, anti-aligned cliques had greater task decodability than aligned cliques. d) The strongest (highest and lowest) anti-aligned and and aligned 20-cliques across the entire cortex. e) The decoding accuracy for the anti-aligned versus aligned ΔNC cliques. f) We evaluated the impact of NCs by destroying correlated variability when training the linear decoder. This was achieved by randomly shuffling task block structure for each brain region separately. g) We computed the difference in decoding performance between unshuffled and shuffled conditions. Shuffling task blocks impacted the decoding performance for anti-aligned cliques significantly more than aligned cliques. This is consistent with the hypothesis that the correlation structure of anti-aligned cliques are important for improved task information decoding (since ΔNCs are reconfigured in the opposite direction of SCs). (*** indicates p*<*0.0001; ** indicates p*<*0.001; * indicates p*<*0.05) Box plot bounds define the first and third quartiles (across participants), box whiskers indicate the 95% confidence interval, and the center line indicates the median.

We implemented this by thresholding the signal-noise differential matrix to include either exclusively aligned or anti-aligned NCs, and then searching for cliques within these thresholded matrices (see Methods). To control for the possibility that identifying cliques would identify brain regions from functionally different networks, we first performed an analysis that identified aligned and anti-aligned ΔNC 5-cliques (cliques with 5 brain regions) for every pair of networks. Identifying both aligned and anti-aligned cliques matched to every network-to-network configuration (e.g., Visual to Somatomotor network), guaranteed that differences in task decoding were not due to decoding cliques from different functional networks. (We also show corresponding results for 8– and 10-cliques; Supplementary Fig. 5).

We directly compared the decoding performance of anti-aligned versus aligned cliques for every pair of networks (Fig. 6b). We found that, while not all pair of networks had a statistically significant difference in decoding performance, 24% of network pairs had a significantly higher decoding performance for anti-aligned versus aligned cliques (13/54 network-matched cliques; two-sided Wilcoxon signed-rank test, Bonferroni-corrected p<0.05). (Note that not all network-network pairs contained anti-aligned and aligned 5-cliques, and so those networks were excluded; see matrix elements colored in white, Fig 6b).) Importantly, and as hypothesized, no aligned clique had a greater decoding accuracy than anti-aligned clique. To obtain a global summary statistic, we computed the average decoding accuracy for all anti-aligned cliques (averaged across all networks) and aligned cliques (Fig. 6c). We found that anti-aligned cliques had a significantly higher decoding performance than aligned cliques (accuracy difference=1.1%, p<10e-06). These findings verify that sets of anti-aligned ΔNCs have improved decodability relative to aligned ΔNCs. Moreover, anti-aligned ΔNCs were overwhelmingly NC reductions (96.4% of all anti-aligned ΔNCs were ΔNC<0). These results highlight three key insights: 1) The impact of tNC should be baselined to spontaneous rNC to infer the impact of NCs on task information decoding; 2) The impact of state-related ΔNCs on task information coding can only be interpreted after knowing the underlying SC; 3) Contrary to prior hypotheses in the neuroimaging literature, NC reductions tend to improve task information coding (rather than inhibit communication) (for review, see Gonzalez-Castillo and Bandettini (2017)).

The above task decoding analysis constrained the comparison of anti-aligned and aligned ΔNCs to a specific network pair. However, it is possible that the signal-noise differentials provide useful information about *which* brain regions are involved in optimizing for task information coding. We therefore lifted the constraint of comparing decoding performance between regions within the same networks. Instead, we sought to identify which brain regions are most/least important for task information decoding, by identifying cliques with the strongest anti-aligned/aligned ΔNCs. We identified the 20-clique with the greatest anti-aligned and aligned ΔNCs, as determined by the magnitude of the signal-noise differential. We found that regions with aligned ΔNCs were primarily located in medial prefrontal and posterior cingulate areas (Fig. 6d). See also Supplementary Fig. 5i,j for a map containing all possible aligned and anti-aligned 20-cliques.) This was consistent with earlier results, which found that the DMN had disproportionate number of regions with aligned ΔNCs (Fig. 4h). Importantly, when we computed the decoding performance of the aligned 20-clique, it exhibited a significantly lower decoding accuracy than the anti-aligned 20-clique (accuracy difference=18.24%, p<10e-06; Fig. 6e). (We replicated this finding using whole-cortex 15-cliques and 25-cliques; Supplementary Fig. 5.) This again provides additional evidence that regions with aligned ΔNCs limit task information decoding, while anti-aligned ΔNCs enhance task information coding. Neuroscientifically, these findings also suggest that NCs with the DMN (which are primarily NC increases) inhibit the fidelity of task information coding.

### Destroying task-state correlations impacts the decodability of task information

Supported by theory, our empirical results demonstrate that the alignment of ΔNCs with their underlying SCs impacts task information decoding. However, signal-noise differentials are determined by the relationship of how NCs emerge given the underlying SC. While SC patterns are an intrinsic property of a system (and likely reflect underlying anatomical connectivity; Passingham et al. (2002)), NC is reflected in ongoing, block-to-block (or trial-to-trial) activity. Thus we sought to assess if destroying the NCs between brain regions (by shuffling block structure) would impact task decoding.

To destroy the correlated activity between brain regions, we shuffled the block ordering for each brain region separately (Fig. 6f; see Methods). This removed the effect of tNCs when training a decoder. (Note that in the context of a decoding analysis, shuffling happened on the training set within every cross-validation fold to ensure no leakage between train and test sets; see Methods). We computed the decoder accuracy after removing the NCs for both anti-aligned and aligned cliques. When comparing the difference between unshuffled and shuffled decoder performance (Δ_*shu f fle*_), we found that destroying NC structure of anti-aligned cliques significantly reduced its decoding performance (unshuffled accuracy=33.6%; shuffled accuracy=25.3%; p<1e-6; Fig. 6g). While shuffling the tNC for aligned cliques also reduced its decoding performance (unshuffled accuracy=15.4%; shuffled accuracy=11.9%; p<1e-5; Fig. 6g)), removing the effect of NCs had a significantly greater impact on the anti-aligned ΔNCs (Δ_*shu f fle*_ anti-aligned=8.4%; Δ_*shu f fled*_ aligned=3.4%; p<1e-5; Fig. 6g). These empirical findings are consistent with the hypothesis that tNC of anti-aligned cliques significantly enhance information coding relative to aligned cliques, and demonstrate the information-coding relevance of tNC changes.

## Discussion

We leveraged insights from neural coding to interpret large-scale task-state correlation changes in human fMRI data. We first characterized the SCs and NCs of human fMRI data using a multi-task dataset with 26 cognitive tasks, finding that SCs had greater network modularity and segregation than the commonly-used rNC matrix. This suggested that SC may have greater utility than rNC in identifying functional specialization across cortical regions. Next, we sought to understand how NCs emerge from underlying network dynamics. We constructed a linear statistical model to gain an intuition of how state-dependent NCs interact with each other. This model revealed that – under the assumption of linear dynamics – tNC should emerge as the algebraic sum of spontaneous background activity and stimulus-specific activity. This implied that stimulus-specific NCs should always align with the underlying SC. In contrast to this model, we did not find this pattern in empirical NCs. Instead, a majority of ΔNCs were anti-aligned with the underlying SCs. This led us to interpret these anti-aligned ΔNCs through a neural coding perspective, which predicts that anti-aligned ΔNCs should improve task information coding. This is because the NCs are reconfigured to avoid interference along the SC axis. Indeed, when testing this prediction in empirical data, we found that anti-aligned ΔNCs had significantly higher task decoding accuracies than ΔNCs that were aligned with their underlying SCs. Together, these findings provide a task information coding perspective to interpret task-state correlation changes in human functional brain networks.

In the human neuroimaging literature, studies of inter-region communication are viewed through the lens of “functional connectivity”. While FC is a broad umbrella term that incorporates a variety of techniques (Reid et al., 2019; Cliff et al., 2022; Frässle et al., 2018; Friston, 2011; Sanchez-Romero and Cole, 2021), the most commonly-used measure is the Pearson correlation – the same metric used in computing spike count NCs. Yet despite the use of identical statistical metrics across the human neuroimaging and neurophysiology, the frameworks for interpreting correlations diverge. On one hand, human neuroimaging studies often analogize the strength of correlation with the strength of “communication” (for review of the literature, see Gonzalez-Castillo and Bandettini (2017)). On the other hand, NCs are typically viewed through the lens of how they impact task information coding (Abbott and Dayan, 1999; Panzeri et al., 2022; Cohen and Kohn, 2011). Empirically, we found that the majority of NCs that enhance information coding – anti-aligned ΔNCs – tend to be tNC decreases (96.4% of anti-aligned ΔNCs are decreases). This finding places these two views at odds, since prior interpretations of tNC reductions have been interpreted as “reduced” or segregated communication among brain regions (Wig, 2017; Rubinov and Sporns, 2010). Here we suggest that the neural coding perspective provides a parsimonious explanation for why reduced tNCs are widespread and enhance task coding: The anti-alignment of the ΔNC with the SC minimizes the amount of signal interference between the two brain regions. Given that the majority of brain regions have a positive SC (Fig. 2b), it follows that the majority of ΔNCs should be reductions to enhance task information coding among brain regions.

There are several conceptual and methodological differences in neural correlation studies between fMRI and electrophysiology literatures relevant to our study. First is the use of the term noise. In neural spiking data (particularly after spike sorting), the term neural noise implies neural activity that cannot be accounted for by experimenter-controlled parameters (e.g., spontaneous activity). In fMRI, noise can emerge from a multitude of sources in addition to spontaneous activity, such as physiological noise, thermal noise, and participant motion. While we employed noise removal methods to triangulate the neural activity of interest (i.e., spontaneous neural activity) (Ciric et al., 2017), it is possible that the effect of the noise correlations we measured were not necessarily purely neural in nature. Nevertheless, it will be important for future work – such as approaches that use simultaneous fMRI imaging with neurophysiology (Shahsavarani et al., 2023; Ma et al., 2016), and those that leverage better individualized estimation methods for fMRI activity (Prince et al., 2022) – to directly verify whether the impact of noise correlations on fMRI decoding originates from neural sources. Second, we used resting-state activity to model baseline neural correlations, whereas electrophysiology studies typically use a baseline control condition. This makes it difficult to fully determine whether changes in neural correlations are due to the effects of an unconstrained neural state. It will be pertinent for future work to assess the impact of noise correlation structure during unconstrained and constrained states. Finally, most studies in the fMRI literature typically estimate tNC using adjacent time points during task performance blocks (Cole et al., 2014; Krienen et al., 2014), rather than the across-trial correlations commonly employed in electrophysiology and also in this study. However, computing the correlation across adjacent timepoints within a task block can make it difficult to disambiguate signal and noise sources, if proper removal of the mean task effect is not performed (Cole et al., 2019). The present approach disambiguates SC and NC measurements by isolating the cross-block mean and cross-block variance by obtaining block-to-block activation estimates separately. Importantly, this is the common approach to calculating SCs and NCs in the neurophysiology literature (Cohen and Kohn, 2011). Nevertheless, to demonstrate the generality of the statistical inferences made here, we found a high correspondence between the ΔNC matrices when computing tNC across timepoints during task blocks (Supplementary Fig. 1). (This is the commonly-used approach to estimating tNC in fMRI neuroimaging.) Together, these findings suggest that differences in tNC calculation should not influence the present conclusions.

Our findings are also widely consistent with prior studies across subfields in neuroscience that find widespread decorrelations during task states. These studies revealed that during task and attentional states, correlations are reduced among pairs of neurons (Cohen and Maunsell, 2009), cortical regions accessed with wide-field calcium imaging data (Pinto et al., 2019), mean-field multi-unit recording across cortical regions in non-human primates (Ito et al., 2020), and human fMRI correlations (Ito et al., 2020). While prior literature has demonstrated that spike count correlations impact information coding in non-human primates (Cohen and Maunsell, 2009; Ni et al., 2022), it was an open question as to whether these intuitions would scale to larger spatial organizations and broader cognitive tuning curves. Our findings affirm that the generic statistical principles developed to understand neural coding in spiking units are translatable to different data modalities, and naturally scale up to broader spatial and cognitive levels of organization. However, the current study only takes into account task-general changes to NCs, rather than task-specific NC changes. While prior work in non-human primate spike recordings suggest that NCs change to support task coding *in general* (rather than optimally for each task) (Ni et al., 2022), it will be important for future studies to investigate the contribution of task-general NC changes versus task-specific NCs to support task information coding.

The present findings, as well as current limitations, open new questions that future studies can explore. First, while the finding that anti-aligned correlations improve the task decodability of those brain regions and networks, it is unclear how this optimized information is implemented and used by the brain. Which downstream brain regions read-out this information? What are the biophysical mechanisms that produce anti-aligned ΔNCs? Future work can build on this work to investigate how optimized task information is used and implemented by the brain (De-Wit et al., 2016). Second, the intuitions behind how the signal-noise differential impacts task coding were developed for two dimensions (i.e., two regions or neurons) (Fig. 5) (Moreno-Bote et al., 2014; da Silveira and Berry, 2014). While we demonstrate that these intuitions generally apply for more than just two regions (e.g., improved decoding for anti-aligned *n*-cliques), it is not explicitly clear how these intuitions generalize to greater dimensions. It will be important for future work to develop theory and measures (beyond just the signal-noise differential/angle) beyond two dimensions. Third, there is a rich fMRI literature on time-varying FC and dynamic co-activation patterns at rest (Lurie et al., 2020; Hutchison et al., 2013). However, in our analysis, we assumed rNC to be stable, as other studies have shown static and dynamic FC represent largely similar information (Matkovič et al., 2023). Nevertheless, it will be important for future studies to assess the impact of large-scale spontaneous dynamics and how they might influence neural coding. Finally, interpreting the impact of NCs on task coding requires knowledge of the underlying SC. In many cases and existing datasets, however, identifying the SC is infeasible, since it requires many tasks and conditions. It will be interesting for future work to develop techniques to approximate the SC without acquisition of task data, such as anatomical connectivity fingerprinting, which has been thought to define the functional tuning of local brain regions (Passingham et al., 2002).

In conclusion, we disambiguate SC and NC in large-scale human functional brain networks using a multi-task fMRI dataset, and characterize the impact of NCs on task information coding. This work bridges the disparate fields of the spike count correlation analyses (typically carried out in non-human animals) with the emerging field of task-state functional connectomics in humans. Importantly, our findings place functional connectomics within a broader framework of neural coding, demonstrating the impact of task-state FC for task coding. We hope these findings spur future investigations into understanding the properties of task information coding in large-scale human brain networks.

## Methods

### Multi-domain task battery dataset

Portions of this section are paraphrased from the dataset’s original publication’s Methods section (King et al., 2019), and a prior study we used to investigate multi-task cortical representations (Ito and Murray, 2023).

We used the publicly available multi-domain task battery (MDTB) dataset, which was originally published to study the functional (task) boundaries of the human cerebellum (King et al., 2019). The dataset contains both resting-state and task-state fMRI data for 24 subjects collected at Western University (16 women, 8 men; mean age = 23.8 years, s.d. = 2.6; all right-handed; see King et al. (2019) for exclusion criteria). All participants gave informed consent under an experimental protocol approved by the institutional review board at Western University.

The MDTB dataset collected data during 26 cognitive tasks, and up to 45 different task conditions for each participant. Tasks were grouped together in two sets (set A and B; Fig. 1e). Each participant first performed all tasks in set A, and returned for a second session to perform tasks in set B. Each task set consisted of two imaging runs. Half of the subjects had sessions separated by 2-3 weeks, while the other half had sessions separated by roughly a year. A separate resting-state scan with two 10 minute runs each was collected for 18/24 subjects. (This resting-state scan was independent of the ‘rest’ block in the task imaging sessions.)

The MDTB dataset was designed to target diverse cognitive processes. Set A and B contained eight overlapping tasks (e.g., theory of mind and motor sequence tasks), and nine tasks unique to each set (Fig. 1a). Both sets contained 17 tasks each. Further details about the experimental tasks and conditions have been previously reported in the original dataset publication (see Supplementary Table 1 of King et al. (2019); https://static-content.springer.com/esm/art%3A10.1038%2Fs41593-019-0436-x/MediaObjects/41593_2019_436_MOESM1_ESM.pdf).

Tasks were performed once per imaging session. Tasks were presented in an interleaved block design. Task blocks began with a 5s instruction screen, followed by 30s of continuous task performance. 11 out of 26 tasks were passive and required no motor response (e.g., movie watching). Tasks that required motor responses were made with either left, right, or both hands using a four-button box using either index or middle fingers. All tasks (within each set) were performed within a single imaging run, ensuring a common baseline between tasks for all participants.

### fMRI preprocessing

Portions of this section are paraphrased from a prior study using a similar preprocessing strategy (Ito and Murray, 2023). fMRI data were minimally preprocessed using the Human Connectome Project (HCP) preprocessing pipeline. The HCP pipelines were implemented within the Quantitative Neuroimaging Environment & Toolbox (QuNex, version 0.61.17; Ji et al. (2022)). The HCP preprocessing pipeline consisted of anatomical reconstruction and segmentation, EPI reconstruction and segmentation, spatial normalization to the MNI152 template, and motion correction.

### fMRI task activation estimation

Portions of this section are paraphrased from a prior study using a similar preprocessing strategy (Ito and Murray, 2023).

We performed a single-subject beta series regression (Rissman et al., 2004) on fMRI task data to estimate parcel-wise activations using the Glasser atlas (Glasser et al., 2016). Each task block (30s) was modeled with a separate task regressor. The instruction period prior to the task block was not included. Thus, the number of task regressors was equivalent to the total number of task blocks per imaging session. Each task regressor was modeled as a boxcar function from the block onset to offset (0s indicate off, 1s indicate on), and then convolved with the SPM canonical hemodynamic response function to account for hemodynamic lags (Friston et al., 1994). We used the coefficients of each regressor as the activation for each task block. Task GLMs were implemented in python using the LinearRegression function within scikit-learn (version 0.23.2) in Python (version 3.8.5). Task GLMs were performed simultaneously with nuisance regression. This was done in an effort to isolate non-neural noise sources when estimating task activations and spontaneous activity. Importantly, the task was modeled simultaneously with noise parameters, as prior studies have shown that removing noise parameters in sequence can artificially induce artifacts (Lindquist et al., 2019). The noise parameters we removed were six motion parameters, their derivatives, and the quadratics of those parameters (24 motion regressors in total). We removed the mean physiological time series extracted from the white matter and ventricle voxels, their derivatives, and the quadratics of those time series (8 physiological nuisance signals). In total, there were 32 nuisance regressors. For task fMRI data, nuisance regressors were included simultaneously with task regressors to extract the task activation estimates described below.

### SC and NC estimation

The SC between two brain regions was computed through the following steps. The mean activation of each task was computed by averaging the block-wise GLM coefficients for that task. This resulted in 26 task activations for every brain region. The SC was then computed as the across-task correlation. Note that since 8 of the 26 tasks were performed in both task sets (i.e., set A and set B), we only included data from one of the task sets. This helped to control the data imbalance across different tasks, ensuring that every task had an equal number of blocks when calculating the mean task activation (16 blocks).

The NC for two brain regions was estimated for each task separately. NC estimation that we performed is identical to task-state FC calculation using a beta series regression (Rissman et al., 2004).

To estimate tNC using task blocks, block by block activation coefficients were obtained for each task separately. Each task had 16 blocks across all imaging sessions. The NC for a pair of regions was the across-block correlation *within* a task. Since there were 26 tasks, there were 26 NCs for every pair of brain regions. We averaged the NC across all tasks (excluding the rNC) to obtain a task-general tNC matrix. rNC was computed using the resting-state blocks during the task imaging session. To verify that the beta series regression approach to calculating NC is similar to other NC calculation approaches (i.e., using timepoints within a task block), we also compared NC estimates using correlations across timepoints (Supplementary Fig. 1). Importantly, rNCs, tNCs, and ΔNCs were highly similar to each other despite differences in how they were estimated (block-to-block activity versus timepoint to timepoint activity). This indicated that task coding properties of NCs evaluated here generalize to both block-to-block and timepoint-to-timepoint NC estimates.

Note that timepoint-to-timepoint NC estimation performed in Supplementary Fig. 1 is consistent with prior approaches to calculating NC (Cole et al., 2019; Ito et al., 2020). Specifically, for each task, we fit a finite impulse response model (across blocks of the same task) to remove the mean-evoked response (which includes the hemodynamic response). This approach flexibly removes the mean-evoked response, while taking into account each brain region’s idiosyncratic hemodynamic response shape. We removed hemodynamic effects of up to 20 seconds after task block offset. This ensured that task signal from the previous block would not overlap (interfere) with task signals in the next block. NCs were then calculated on the residual time series. This approach ensured that NCs were not conflated by the mean (i.e., signal) response. We also did not directly compare tNC with rNC using data from the separate resting-state fMRI scan. This was because the data obtained during the task sessions spanned over 5 hours, whereas the separate resting-state scan was only 10 minutes long. This made it impossible to perform a direct comparison of task-to rest-state fMRI using the same exact temporal intervals in this dataset.

### Network analysis

We performed both network-style (Rubinov and Sporns, 2011) and gradient-style (Huntenburg et al., 2018) analysis on SC and NC matrices. Network-style analysis included computing the network modularity and network segregation of SC and rNC matrices with respect to a previously-published functional network partition (Ji et al., 2019).

We used an undirected signed modularity metric that calculates modularity with respect to a provided network partition (Rubinov and Sporns, 2011). We use the asymmetric variant that treats positive and negative values differently (i.e., positive values link nodes within a module, and negative values dissociate nodes between modules). Modularity was calculated as

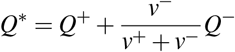

where *Q*^±^ is defined as

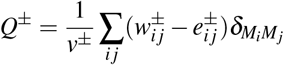

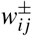 is the connection weight (positive or negative values only), 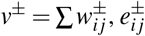 is the chance-expected within module connections defined as 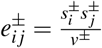, where 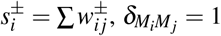 when *i* and *j* are in the same network module and 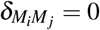 otherwise. Code was implemented using the brain connectivity toolbox (bctpy version 0.5.0).

Network segregation was measured as the difference between within-module and between-module weights, divided by within-module weights (Chan et al., 2014). Segregation was first calculated for each region separately, and then averaged across all regions. Segregation of a region *i* was computed as

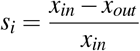

where *x*_*in*_ is the within-module weights for region *i*, and *x*_*out*_ is the between-module weights.

Gradient-style analysis was computed by performing a Principal Components Analysis (PCA) on either the SC or NC matrices. SC and NC matrices were thresholded to retain only 20% of the strongest correlations prior to calculating gradients. PCA was implemented using scikit learn’s PCA function (sklearn.decomposition.PCA, version 1.0.2).

### Linear statistical network model

We used a statistical model to predict how task-related variability influences baseline spontaneous activity. We partitioned 300 nodes into 10 networks (30 nodes each). Networks were fully connected with a weight of 1. Since resting-state NC typically exhibits positive correlations among all pairs of regions, we introduced a globally shared signal. Specifically, a region’s activity *x*_*i*_ was determined as the linear sum of four Gaussian distributions (10,000 samples), *X ∼ N*(0, 1):

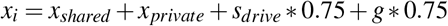

where *x*_*shared*_ is a shared time series amongst nodes within a network, *x*_*private*_ is a unique time series for *x*_*i*_, *s*_*drive*_ is the stimulus-related variance (set to 0 in the baseline spontaneous case), and *g* is the global variance shared by all nodes.

### Task decoding analyses

#### Clique identification

We identified cliques of aligned and anti-aligned signal-noise differentials. This was to test the impact of signal-noise differentials on task information decoding. Cliques are a sub-network of a graph that are fully connected (Sizemore et al., 2018). This means that every node is connected to every other node in that sub-network. Using the signal-noise differential matrix (Fig. 6a), we identified aligned and anti-aligned cliques by creating thresholded matrices of exclusively aligned and anti-aligned region pairs, respectively. This ensured that when performing decoding analyses on a set of regions, that every pair of region was either aligned or anti-aligned. This was important given that, from a neural coding perspective, aligned signal-noise differentials would be information-limiting relative to baseline correlations, while anti-aligned signal-noise differentials would be information-enhancing (Fig. 5).

We identified aligned and anti-aligned 5-cliques for every pair of functional network configuration (e.g., region sets between the Default Mode with the Frontoparietal network; Visual 1 network to the Somatomotornetwork). Matching aligned and anti-aligned 5-cliques to the specific network configuration controlled for the possibility of inherent differences in decoding performance when identifying cliques from different networks. (For example, there might be intrinsic differences when comparing the decoding performance of an aligned clique in the Default Mode network versus the Visual 1 network.) However, aligned and anti-aligned 5-cliques did not exist for all network pairs. These network pairs were therefore excluded from analysis, since aligned and anti-aligned decoding performances could not be directly compared. In supplementary analyses, we also show that our findings generalize to 8-cliques and 10-cliques (Supplementary Fig. 5).

In addition, we identified the 20-clique with the strongest aligned and anti-aligned 20-clique (Fig. 6d). Strongest was defined as having the greatest negative or positive average values within an aligned or anti-aligned 20-clique (using the signal-noise differential matrix; Fig. 6a). Note that while the aligned and anti-aligned 20-cliques were in spatially disjoint regions, decoding performance was appropriately controlled for by removing the effect of correlated variability (e.g., Leavitt et al. (2017); Fig. 6f,g), as discussed in the next subsection.

Clique identification was carried out using the python package NetworkX (networkx.find_clique function; version 2.5). To make identifying cliques more tractable, the signal-noise differential matrix was thresholded to retain only the top 20% positive or negative values. Note that we also replicated these findings using 15– and 25-cliques (Supplementary Fig. 5). For the 25-clique analysis, we thresholded the signal-noise differential matrix to the top 40% of positive or negative values.

#### Decoding analyses

To assess the role of NCs on task information coding, we performed a multi-task (26-way) linear decoding analyses. Decoding analyses were performed within subjects, using the block-wise activations of every task. There were 26 tasks with 16 blocks per task (416 samples per subject). We performed a leave-one-out cross-validation, cross-validating across blocks. Samples in the training set were bootstrapped (20 samples per task type, with replacement). Prior to fitting the linear decoder on the training sets, samples in the training set were feature-normalized (z-normalized), and samples in the test set were also feature-normalized using the mean and standard deviation estimated from the training set (to avoid train-test leakage). A linear decoder was fit using logistic regression, and was implemented using scikit learn (version 1.0.2).

To evaluate the effect of correlated variability on aligned and anti-aligned cliques, we performed a follow-up analysis that removed the impact of NCs on linear decoding. This was implemented by shuffling the ordering of task blocks for each brain region and each task type separately (see Fig. 6f). This was done on the training set within each cross-validation fold. Shuffling blocks for each brain region separately removed the contribution of NCs on training a linear decoder.

Note that while prior studies suggest that the best practices for decoding analyses employ a 5– or 10-fold cross-validation (Varoquaux, 2018), we used a leave-one-out cross-validation approach to maximally assess the impact of correlated activity (within the training set) on decoding performance. (Note exactly one sample from each task was left out from the training set, such that the test set had 26 samples in total.) Moreover, we were not focused on making inferences on decoding performance relative to chance. Instead, we were interested in assessing how correlated activity (within the training set) impacted decoding performance for aligned and anti-aligned cliques, and how shuffling correlated activity would (within the training set) would impact overall decoding performance. If NCs had no impact on decoding performance, shuffling the block structure in the training set would have no impact on task decoding.

## Data visualization

All graphical plots were visualized using seaborn (version 0.11.2; Waskom (2021)). All cortical surface plots were visualized using surfplot (version 0.1.0; Gale et al. (2021); Vos de Wael et al. (2020)).

## Code and data availability

All data in this study has been made publicly available on OpenNeuro by King and colleagues (King et al., 2019). https://openneuro.org/datasets/ds002105

All code related to this study will be made publicly available on GitHub. Analyses and models were implemented using Python (version 3.8.5).

## Acknowledgements

This project was supported by NIH grant R01MH112746 (JDM), NSF NeuroNex grant 2015276 (JDM), and a Swartz Foundation Fellowship (TI). The authors acknowledge the Yale Center for Research Computing at Yale University for providing access to the Grace cluster and associated research computing resources.

## Supplementary Figures

**Supplementary Figure 1.**
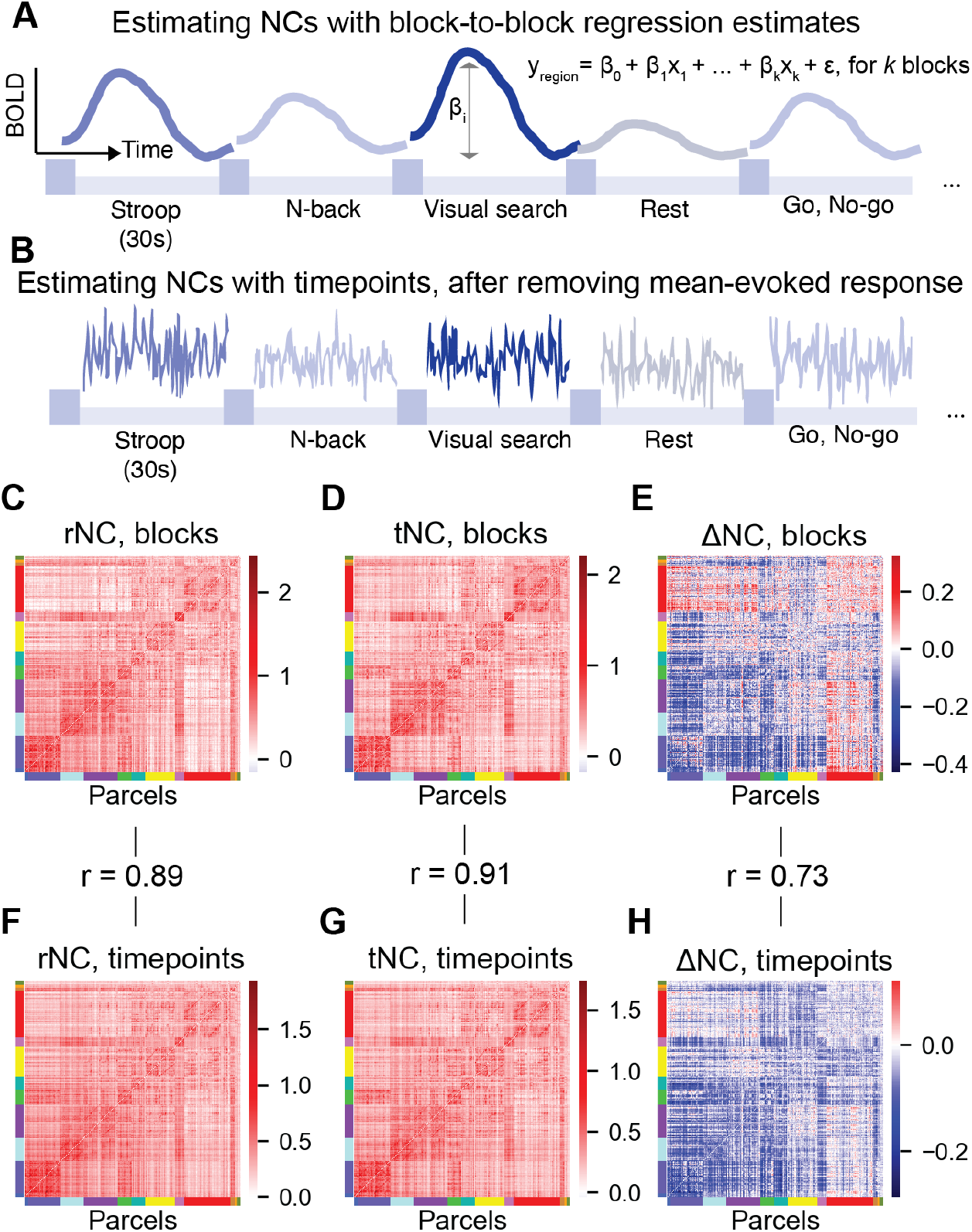
NCs computed using block-to-block activity estimates versus timepoint-to-timepoint estimates reveal quantitatively similar NC estimates. a) To estimate NCs using block-to-block estimates, we performed a beta series regression. In a beta series regression, every block (or trial) has its own independent regressor. Every block/trial therefore has its own activity estimate. (Image is a schematic.) b) To compare NCs using the more traditional approach, we estimated NCs using correlations estimated across timepoints within task blocks. To ensure task-driven variance/noise was not conflated with the mean-evoked (i.e., signal) response, we performed a finite impulse response model across all blocks for each task type separately (Cole et al., 2019). This ensured that NCs were computed using the background task-driven variance. c) The rNC matrix computed using rest blocks (as implemented in the main text). d) The tNC matrix computed using task blocks, averaged across all tasks (as implemented in the main text). e) The ΔNC matrix using block-wise NC estimates (as implemented in the main text). f) The rNC matrix computed as the correlation across timepoints. Resting-state blocks were first concatenated across all imaging sessions for a participant. The rNC was then computed on the concatenated time series. g) The tNC matrix computed as the correlation across timepoints. Blocks for each unique task were first concatenated for each participant. tNC was computed for each task, and then averaged across all tasks to obtain a task-general NC matrix. h) The ΔNC matrix using timepoint-to-timepoint NC estimates. Despite being computed using different approaches (with varying amounts of data per NC), rNCs (r=0.89), tNCs (r=0.91), and ΔNCs (r=0.73) were highly similar across these approaches.

**Supplementary Figure 2.**
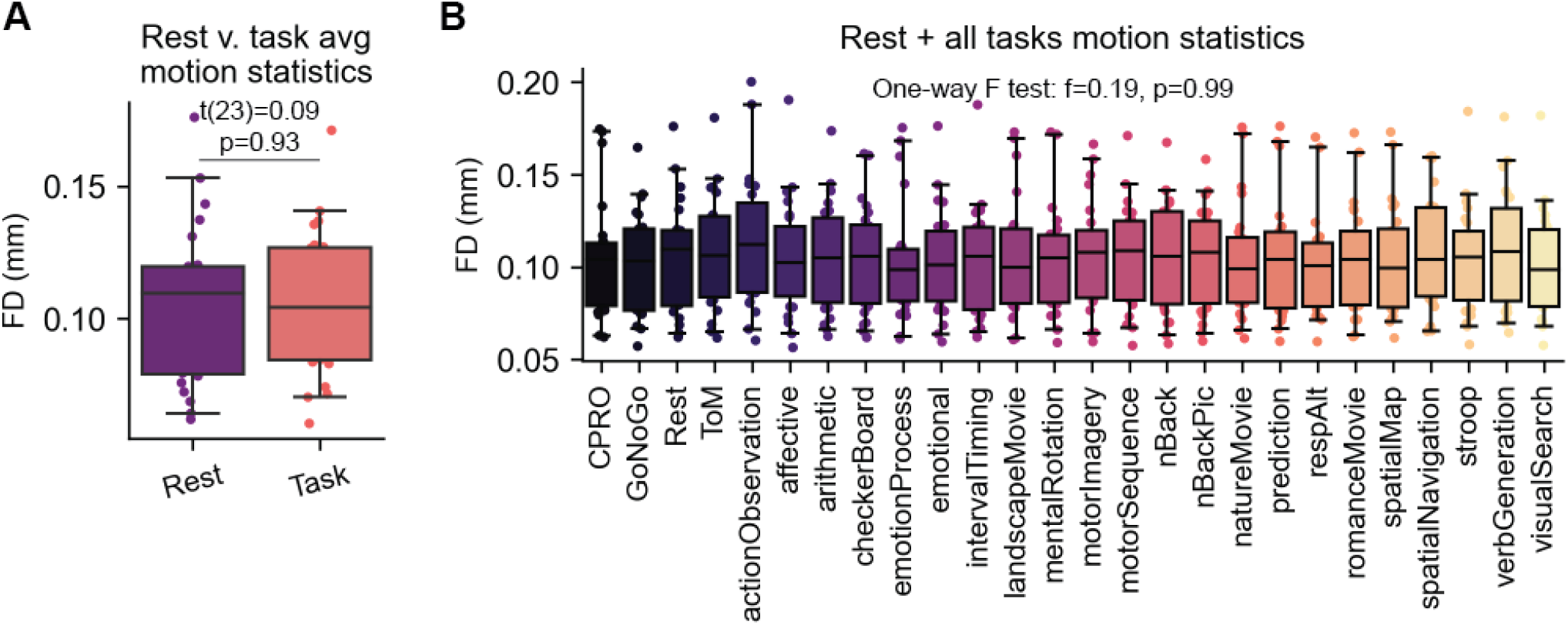
Motion statistics for each subject across rest and task states. a) We computed the framewise displacement (FD; Power et al. (2012)) for all timepoints during rest and task blocks. We found no significant difference between the average FD between rest and task states. b) The average FD during the blocks across rest and each task individually. We performed a one-way F test to assess if the FD of a given state was statistically different from the FD of all other states. We found no statistically significant deviation of FD across all states. Box plot bounds define the first and third quartiles of the (across participant) distribution, box whiskers indicate the 95% confidence interval, and the center line indicates the median.

**Supplementary Figure 3.**
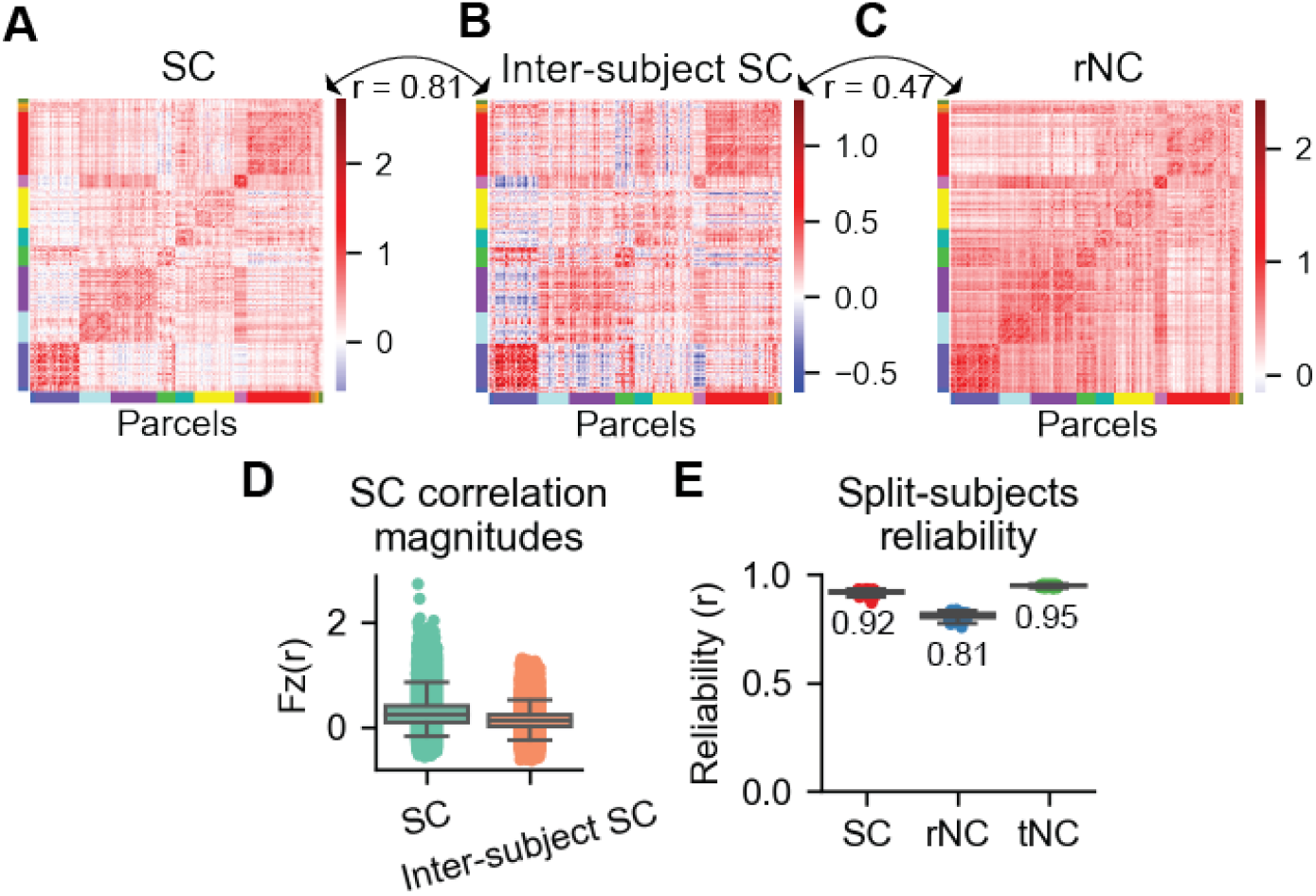
Reliability of the SC, rNC, and tNC matrices. a) The group-averaged SC matrix (same as in the main text). b) The inter-subject SC matrix. The inter-subject SC removes the potential confound of noise sources that are idiosyncratic to the participant and/or scan session (Kim et al., 2018). This was calculated by taking one subject’s multi-task activation vector for a single brain region, and then correlating that vector with the group-averaged (excluding that one subject) multi-task activation vector from all other brain regions. This approach exclusively captures SC at the group-level. We found that the group-averaged SC matrix and the inter-subject SC matrix maintained a high correspondence (r=0.81), suggesting that in general, the organization of the group-averaged SC captures the organization of signal correlations that are not subject-specific. c) The group-averaged rNC matrix (same as in main text), for comparison. d) The distribution of correlation values between the group-averaged SC (a) is wider than the distribution of the inter-subject SC (b). This is because inter-subject SC removes sources of correlated variability that are idiosyncratic to an individual. Boxplots reflect the distribution across all pairs of correlations. Box plot bounds define the first and third quartiles, box whiskers indicate the 95% confidence interval, and the center line indicates the median. e) Group-averaged SC, rNC, and tNC measurements are overall reliable. We performed a splits subject analysis, randomly sampling half the subjects SC/rNC/tNC, and correlating it with the SC/rNC/tNC of the other half, respectively. Across participant splits, SC, rNC, and tNC reliability was above r=0.81. rNC had the lowest reliability, which is likely due to the fact that rNC correlations were computed using only 16 samples per pair of parcels. (tNC is computed by correlating 16 samples per pair of parcels per task, and then averaged across tasks. SC is computed by estimating the average activation for each task, and then correlating across 25 tasks.) We bootstrapped 100 random splits; box plot bounds define the first and third quartiles of those 100 splits, box whiskers indicate the 95% confidence interval, and the center line indicates the median.

**Supplementary Figure 4.**
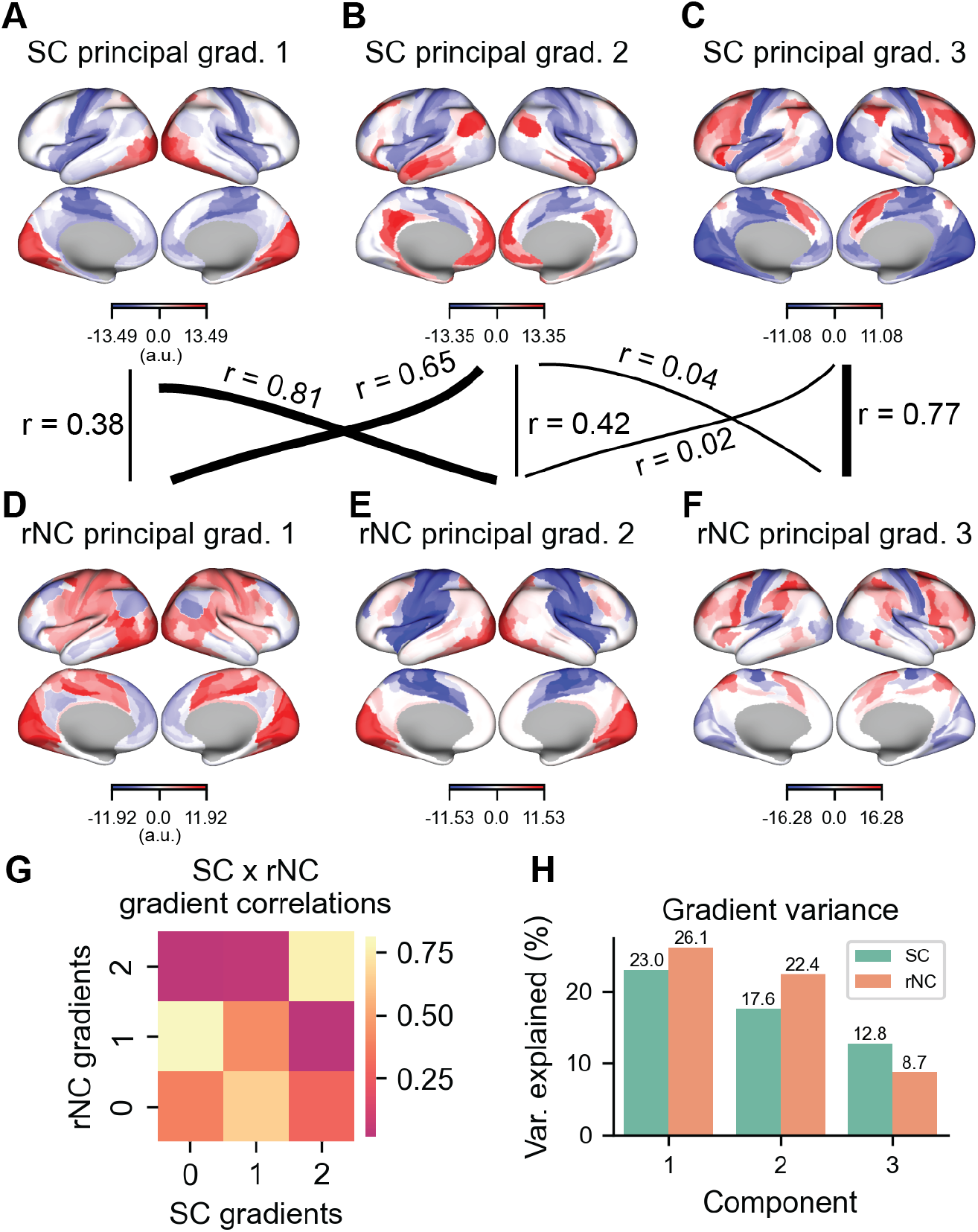
Detailed comparison of the first three SC and rNC gradients. a) The first, b) second, and c) third SC gradient. d) The first rNC gradient, which has highest similarity to the second SC gradient. e) The second rNC gradient, which has highest similarity to the first SC gradient. f) The third rNC gradient, which has greatest similarity to the third SC gradient. Together, these findings suggest that the first three dimensions of SC and rNC are similar, but that the first two components are flipped in SC and rNC. Note that correlation values reflect the absolute value, since the orientation of PCA loadings are arbitrary. g) All pairwise correlations (absolute value) between the first three SC and rNC gradients. h) The variance explained of each gradient (principal component) for each SC and rNC matrix.

**Supplementary Figure 5.**
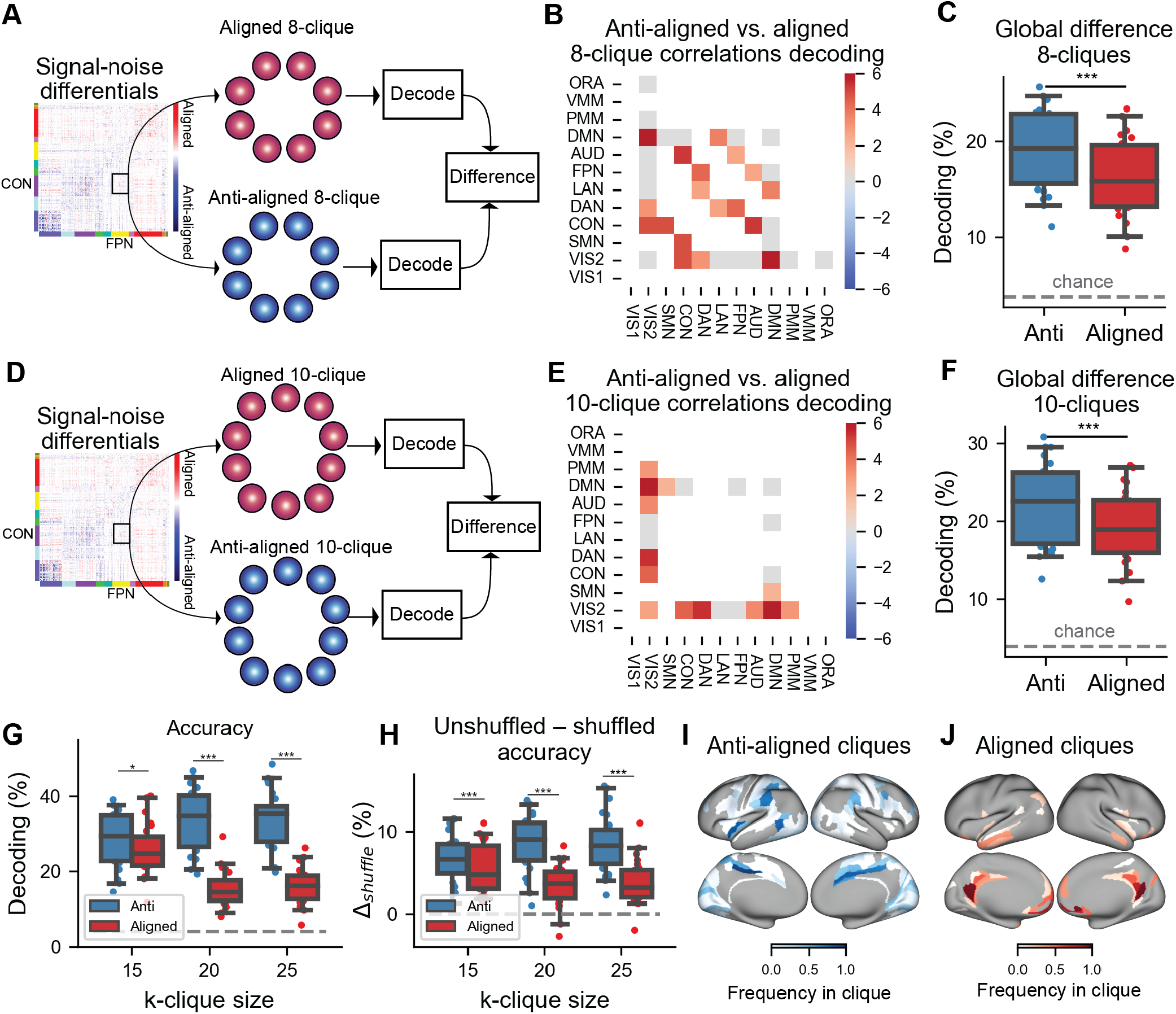
Decoding analyses for different k-clique sizes – supplementary analyses for Fig. 6. a) Identifying 8-cliques within every pair of networks. b) Decoding accuracies for anti-aligned versus aligned 8-cliques for every pair of networks. Note that gray matrix elements indicate non-significant differences, and white elements indicate non-testable network configurations (due to non-existence of anti-aligned and/or aligned cliques of that size). c) Anti-aligned versus aligned decoding accuracies, averaged across all available network pairs. d-f) Same as a-c, but using 10-cliques. g) Decoding accuracies for 15, 20 (in the main text), and 25 anti-aligned and aligned cliques identified across the entire cortex. Anti-aligned cliques consistently had greater decoding accuracies than aligned cliques. h) The difference between unshuffled and shuffled decoding accuracies for anti-aligned and aligned cliques. Removing NCs impacted anti-aligned cliques significantly more than aligned cliques. i) We identified all possible 20-cliques for anti-aligned and j) aligned NCs, and plotted the frequency with which each region appeared in all cliques. Anti-aligned cliques tended to reside in sensory and motor areas, awhile aligned cliques were most frequently observed in medial prefrontal and posterior cingulate areas. (*** indicates p<0.0001; ** indicates p<0.001; * indicates p<0.05)

